# Riluzole does not ameliorate disease caused by cytoplasmic TDP-43 in a mouse model of amyotrophic lateral sclerosis

**DOI:** 10.1101/749846

**Authors:** Amanda L. Wright, Paul A. Della Gatta, Sheng Le, Britt A. Berning, Prachi Mehta, Kelly R. Jacobs, Hossai Gul, Rebecca San Gil, Thomas J. Hedl, Winonah R. Riddell, Owen Watson, Sean S. Keating, Juliana Venturato, Roger S. Chung, Julie D. Atkin, Albert Lee, Bingyang Shi, Catherine A. Blizzard, Marco Morsch, Adam K. Walker

**Author notes:** Correspondence should be addressed to: Dr Adam K. Walker.

## Abstract

Amyotrophic lateral sclerosis (ALS) is a neurodegenerative disease commonly treated with riluzole, a small molecule that may act via modulation of glutamatergic neurotransmission. However, riluzole only modestly extends lifespan for people living with ALS and its precise mechanisms of action remain unclear. Most ALS cases are characterised by accumulation of cytoplasmic TAR DNA binding protein of 43 kDa (TDP-43), and understanding the effects of riluzole in models that closely recapitulate TDP-43 pathology may provide insights for development of improved therapeutics. We therefore investigated the effects of riluzole in transgenic mice that inducibly express nuclear localisation sequence (NLS)-deficient human TDP-43 in neurons (*NEFH*-tTA/*tetO*-hTDP-43ΔNLS, ‘rNLS’, mice). Riluzole treatment from the first day of hTDP-43ΔNLS expression did not alter disease onset, weight loss or performance on multiple motor behavioural tasks. Riluzole treatment also did not alter TDP-43 protein levels, solubility or phosphorylation. Although we identified a significant decrease in GluA2 and GluA3 proteins in the cortex of rNLS mice, riluzole did not ameliorate this disease-associated molecular phenotype. Likewise, riluzole did not alter the disease-associated atrophy of hindlimb muscle in rNLS mice. Finally, riluzole treatment beginning after disease onset in rNLS mice similarly had no effect on progression of late-stage disease or animal survival. Together, we demonstrate specific glutamatergic receptor alterations and muscle fibre-type changes reminiscent of ALS in rNLS mice, but riluzole had no effect on these or any other disease phenotypes. Future targeting of pathways directly related to accumulation of TDP-43 pathology may be needed to develop better treatments for ALS.

**Significance Statement:** Accumulation of cytoplasmic TDP-43 protein is the hallmark pathology of ALS. Riluzole is the most widely used drug for ALS treatment, but provides only a short extension of lifespan. We demonstrate here in the rNLS mouse model, which mimics TDP-43 pathology, that riluzole does not ameliorate progressive alterations in motor strength and coordination, muscle atrophy, glutamate receptor levels, or TDP-43 protein levels and solubility, and does not prolong animal survival. Riluzole similarly did not affect decreased levels of glutamate receptor subunits GluA2/GluA3 in rNLS mice. The inability of riluzole to rescue pathological or phenotypic changes in this TDP-43 model provides further impetus for the discovery of improved therapies targeting the key drivers of ALS pathogenesis.

## Introduction

Amyotrophic lateral sclerosis (ALS) is a debilitating and inevitably fatal neurodegenerative disease which causes paralysis due to progressive loss of upper and lower motor neurons of the brain and spinal cord (Robberecht and Philips, 2013). The small molecule 2-amino-6(trifluoromethoxy)benzothiazole, known generically as riluzole, is the most widely prescribed and until recently was the only approved drug therapy for the treatment of ALS (Lacomblez et al., 1996; Fang et al., 2018). However, the efficacy of riluzole is limited, resulting in a short extension in lifespan for people with ALS of only a few months, with little if any individual improvement in quality of life (Lacomblez et al., 1996; Miller et al., 2012). A better understanding of the biological mechanisms driving the onset and progression of ALS is needed, and determining the pathways through which riluzole exerts its modest effects may also aid the development of more effective therapeutics.

The mechanisms of action of riluzole remain unclear despite decades of use in people following FDA approval in 1995, although modulation of glutamatergic neurotransmission is likely involved (Bellingham, 2011). Indeed, people with ALS have increased levels of the excitatory neurotransmitter glutamate in cerebrospinal fluid (Rothstein et al., 1990), and altered excitability of motor neurons may be an early feature of disease (Martinez-Silva et al., 2018; Jiang et al., 2019). Amongst a myriad of other potential disease-relevant effects of riluzole, the drug has also been proposed to affect estradiol transport (Klemann et al., 2018), to inhibit kinases potentially involved in ALS pathology (Bissaro et al., 2018), and to decrease activation of the heat shock response related to protein aggregation (Shaw et al., 2018).

Although riluzole was initially shown to have a modest protective effect in the most widely used mouse model of ALS, the SOD1^G93A^ mouse (Gurney et al., 1996; Gurney et al., 1998), a later more rigorous study demonstrated conclusively that SOD1^G93A^ mice do not benefit from riluzole treatment (Scott et al., 2008). Likewise, in other mouse models of ALS expressing rare ALS-linked mutant TAR DNA binding protein of 43 kDa (TDP-43) or fused in sarcoma (FUS) proteins, riluzole also failed to show any protective effect in function or lifespan (Hogg et al., 2018). Similarly, riluzole has no protective effect in a rat model with ALS-linked mutant TDP-43 expression (Chen et al., 2020). Notably, the hallmark pathology in the vast majority of ALS cases is cytoplasmic accumulation of post-translationally modified wildtype, not mutant, TDP-43 in affected regions of the brain and spinal cord (Neumann et al., 2006; Ling et al., 2013), and none of these previously studied rodent models robustly recapitulate this key pathological feature of ALS. It therefore remained unclear whether riluzole may be effective in a mouse model more closely resembling most cases of human ALS with cytoplasmic TDP-43 pathology.

Here we sought to investigate the therapeutic effects of riluzole in a well-characterised mouse model with cytoplasmic accumulation of TDP-43, the *NEFH*-tTA/*tetO*-hTDP-43ΔNLS (‘rNLS’ mice), which closely recapitulates the key pathology observed in the vast majority of people with ALS (Walker et al., 2015). We found that riluzole was unable to delay onset or alter progression of disease in rNLS mice when drug treatment began prior to symptom onset, by testing the effects of riluzole in early disease. Similarly, we show that riluzole treatment did not affect disease progression or survival of rNLS mice when drug treatment began after functional decline, an approach that more closely mimics the clinical situation of riluzole treatment beginning after symptom onset. Interestingly, we identified a significant decrease in protein levels of the key glutamatergic α-amino-3-hydroxy-5-methyl-4-isoxazolepropinionic acid (AMPA) receptor subunits GluA2 and GluA3 in the cortex of rNLS mice, suggesting disease-related alterations in postsynaptic glutamate signalling. However, riluzole treatment did not affect this loss of GluA2/3 proteins. Likewise, riluzole did not affect TDP-43 protein levels, solubility or phosphorylation, and did not counteract muscle atrophy in rNLS mice. Overall, our findings demonstrate that riluzole has no effect on pathological changes or disease signs in the rNLS mouse model. Our study therefore provides additional emphasis on the need for identification of improved therapeutics for ALS and other neurodegenerative diseases characterised by the presence of TDP-43 pathology.

## Materials and Methods

### Animals

All animal procedures were performed in accordance with the Macquarie University animal care committee’s regulations. Transgenic TDP-43 rNLS mice for expression of human TDP-43ΔNLS were bred under specific pathogen-free conditions at Australian BioResources (Moss Vale, Australia). Monogenic B6;C3-Tg(NEFH-tTA)8Vle/J (*NEFH*-tTA line 8, stock #025397) mice and monogenic B6;C3-Tg(*tetO*-TARDBP*)4Vle/J (tetO-hTDP-43ΔNLS line 4, stock #014650) mice were initially obtained from the Jackson Laboratory (Bar Harbour, ME, USA). Both lines were maintained individually on a mixed B6/C3H background by mating of monogenic male mice of each line to wildtype B6/C3H F1 females, which were obtained by the cross of female C57BL/6JArc mice with male C3H/HeJArc mice (Australian Resources Centre, Canning Vale, Australia). Generally, male *NEFH*-tTA mice were crossed with female *tetO*-hTDP-43ΔNLS mice to obtain bigenic TDP-43 rNLS and monogenic and non-transgenic littermate control mice.

### Animal experiments overview

Mice were transferred to and housed in the central animal facility at Macquarie University for experiments. Mice arrived at least one week prior to beginning of experiments to allow for acclimatisation. For all therapeutic efficacy studies, littermates were used and allocated randomly by personnel independent from authors and experimenters to receive either vehicle or riluzole treatment in a blinded manner. Experimenters were blinded to treatment but not to genotype. A small cohort of male rNLS mice were included in the study initially but were excluded from the dataset due to unforeseen complication of urinary retention in some of the aged male mice. To avoid this confound, all mice outlined in this study were females. All animals were housed under identical conditions in a 12-hour standard light/dark cycle with access to water and food *ad libitum*. rNLS bigenic mice were housed together with matched littermate control mice. All breeding and weaned mice were fed a chow diet containing 200 mg/kg doxycycline (dox, Gordon Specialty Feeds). hTDP-43ΔNLS expression was induced in mice at 6-8 weeks of age by switching all mice to a matched standard chow diet lacking dox (Gordon Specialty Feeds). All mice were supplied with supplementary diet (DietGel 76A, ClearH_2_O) from day 15 after switching to the standard diet to ensure adequate nutrition and hydration as disease progressed.

### Drug administration

Riluzole (2-Amino-6-(trifluoromethoxy)benzothiazole, #R116-500mg, Sigma Aldrich) was dissolved in corn oil (C8267, Sigma Aldrich) to a final concentration of 3 mg/mL. For each treatment, either riluzole (volume calculated based on a dosage of 8 mg/kg) or the equivalent volume of corn oil (vehicle control) were gavaged once per day into the stomach of the mice using a flexible feeding tube (FTP-20-38, Instech Laboratories, Inc). Treatment was started on day 0 (for onset study) or on day 29 (for post-onset study) of switching to the standard diet. All drug treatments were given once daily (between 0900h – 1200h) throughout the experiment until the pre-defined endpoint or until the end-stage of the disease (defined below). All drug and vehicle treatments were coded, administered and analysed by a researcher blinded to the treatment groups.

### Monitoring and behavioural assessments

All mice were weighed and assessed in the morning (between 0900h – 1200h), three times per week as previously described (Walker et al., 2015) from the first day when they were switched to normal diet until the disease end-stage (defined as weight below 80% of initial weight on each of 3 consecutive days, or paralysis of both hindlimbs, or inability to right within 10 seconds from both sides). Motor behaviour was tested once weekly using grip strength, inverted grid and rotarod tests. Three training sessions for each of these tests were performed one week before the switch to standard diet.

*For the neurological score assessment*, mice were scored on a 4 point scale by an individual blinded to treatment group: 0, normal; 1, abnormal hindlimb splay, slightly slower gait, normal righting reflex; 2, hindlimb partially or completely collapsed, foot dragging along cage, slow righting reflex; 3, rigid paralysis in hindlimb or minimal movement, hindlimb is not being used for forward motion, slow righting reflex.

*For the grip strength test*, a digital force meter (Model 47200, Ugo Basile) was used to measure maximal muscle grip strength from all four limbs. Mice were held by the tail and placed in the centre of the grid connected to the force meter. The mice could use all four limbs to grip on the grid. The tail was pulled horizontally to the grid with a smooth steady manner until the mouse released the grid. The peak strength of the grip was measured in gram force. Each mouse was tested 3 times with at least 20 mins resting period between tests. The average of the highest two recordings was taken as the final result for analysis.

*For the inverted grid test of muscle strength (also known as wirehang test)*, mice were placed on a customized metal grid (0.8 cm x 0.8 cm square grid, stainless steel), and gently flipped to be suspended upside-down approximately 40 cm above a lightly padded clean cage. The time to fall was recorded up to a maximum 180 s. If a mouse did not perform the maximum time, three tests were performed with minimal interval of 20 mins resting period between tests, and the average of the longest two performances was taken as the final result for analysis.

*For the rotarod test of motor coordination, balance and stamina*, mice were placed on a rotarod apparatus (Model 7650, Ugo Basile) at a speed of 4 rpm with acceleration up to 40 rpm over 300 s (onset study) or 180 s (post-onset study). The time to fall was recorded, or the maximal time was recorded if mice were still on the rod without falling at the end of a session. Three tests were performed for each mouse with minimal interval of 20 mins, and the average of the longest two performance was taken as the final result for analysis.

### Perfusion and tissue collection

Mice were deeply anesthetized using ketamine (100 mg/kg) and xylazine (10 mg/kg) and transcardially perfused with ∼20 mL room temperature phosphate-buffered saline (0.01M PBS). Brains were removed and divided in half sagittal from the midline. The left hemisphere was quickly dissected to isolate the cortex, and this was immediately frozen on dry ice and stored at -80°C. Tibialis anterior muscles were snap-frozen in liquid nitrogen-cooled isopentane and stored at -80°C.

### Preparation of tissue lysates

Tissues were thawed on ice and then sonicated in 5× v/w RIPA buffer with phosphatase and protease inhibitors (50 mM Tris, 150 mM NaCl, 1% NP-40, 5 mM EDTA, 0.5% sodium deoxycholate, 0.1% SDS, pH 8.0). Samples were centrifuged at 4°C, 100,000 *g* for 30 min and the supernatant taken as the RIPA-soluble fraction. The pellet was washed by sonication with RIPA buffer as above. This supernatant was discarded, and the pellet sonicated in 2× v/w urea buffer (7 M urea, 2 M thiourea, 4 % CHAPS, and 30 mM Tris, pH 8.5) and centrifuged at 22°C, 100,000 *g* for 30 min. This supernatant was taken as the RIPA-insoluble/urea-soluble fraction. Protein concentrations of the RIPA-soluble fractions were determined using the bicinchoninic acid protein assay (Pierce) and volumes of urea-soluble protein used for immunoblotting were normalised to protein content of the matched RIPA sample.

### Immunoblotting

Proteins were resolved by SDS-PAGE electrophoresis on BioRad polyacrylamide gels. For all immunoblots, 20 μg per sample were added to each lane. Electrophoresis was carried out at 120 V for 1.5 hrs. Following electrophoresis, proteins were transferred to nitrocellulose membranes using the BioRad Transblot Turbo system. Membranes were incubated with 3% BSA for 1 hr, followed by exposure to the primary antibody. Primary antibodies included: rabbit anti-human/mouse TDP-43 1:5000 (Proteintech Cat# 10782-2-AP, RRID:AB_615042), mouse anti-human TDP-43 1:5000 (Clone 5104, (Kwong et al., 2014)), mouse anti-phospho-TDP-43 (Ser409/410) (Cosmo Bio Co Cat# CAC-TIP-PTD-M01, RRID:AB_1961900), rabbit anti-EAAT2 1:1000 (Abcam Cat# ab178401), rabbit anti-GluA1 1:1000 (Abcam Cat# ab31232, RRID:AB_2113447), rabbit anti-GluA2 1:1000 (Millipore Cat# AB1768, RRID:AB_2247874), rabbit anti-GluA3 1:1000 (Abcam Cat# ab40845, RRID:AB_776310), and mouse anti-GAPDH 1:5000 (Proteintech Cat# 60004-1-Ig, RRID:AB_2107436), diluted in 0.1% BSA overnight at 4°C. Membranes were washed and incubated with anti-mouse/rabbit LiCor 680/800 secondary antibodies (1:20,000) for 1 hr. Signals were detected using an Odyssey CLx and quantified with ImageStudio software.

### Immunohistochemical analysis of muscle fibre type

Muscle fibre type and cross sectional area were determined as previously described (Bloemberg and Quadrilatero, 2012). Sections at 8 μm of tibialis anterior were affixed to StarFrost Slides (ProScitech, Kirwan, QLD) and frozen at -80°C. For staining, slides were thawed for 10 min at room temperature and blocked in 10% goat serum (Life Technologies, Australia) diluted in PBS (GS/PBS) for 1 h at room temperature. Slides were incubated in primary antibodies diluted in GS/PBS for 2 h at room temperature [BA-F8 (1:20, MHCI), SC-71 (1:50, MHCIIA) and BF-F3 (1:20, MHCIIB) from Developmental Studies Hybridoma Bank; Laminin (1:100) from Sigma Aldrich Cat# L9393]. Sections were washed in PBS and incubated with secondary antibodies [Alexa Fluor goat anti-mouse IgG2b 647, Alexa Fluor goat anti mouse IgG1 488, Alexa Fluor goat anti-mouse IgM 555, Alexa Fluor goat anti-rabbit IgG H+L 405, all 1:500 (Life Technologies)] diluted in GS/PBS for 1 h at room temperature. Slides were washed in PBS and mounted with coverslips using Prolong Gold (Life Technologies). Sections were imaged using an Olympus Fluoview FV10i confocal microscope (Olympus, Australia). All images were acquired using the same laser and sensitivity settings, and 5-8 random representative images of the tibialis anterior were taken at 20x magnification for cross sectional fibre size analysis and fibre typing. Muscle cross sectional area was determined using Olympus CellSens software (Olympus, Australia) via a custom-made macro developed by the authors.

### Statistical analyses

All statistical analysis was performed using the statistical package Prism 7 (Graphpad). For normally distributed data, differences between means were assessed as appropriate, by unpaired two-tailed t-test or two-way ANOVA with or without repeated measures followed by Sidak’s multiple comparisons test. Simple effect tests were performed when group interactions were identified by ANOVA. The *p*-value, and F-statistic from ANOVA tests and associated degrees of freedom (between groups and within groups, respectively) are reported in parentheses. For Kaplan-Meier survival curves, Log-rank (Mantel-Cox) test was conducted. All data is presented as mean ± SEM. For all statistical tests, a *p*-value of <0.05 was considered to be significant.

## Results

We conducted two investigations in the rNLS mouse model of TDP-43 pathology: a pre-disease onset study to determine the effects of riluzole when treatment began prior to motor symptoms, and; a post-disease onset study to investigate the more clinically relevant scenario of riluzole treatment beginning after development of motor deficits.

### Riluzole does not delay onset or progression of motor deficits in rNLS mice

The rNLS mice used in this study express the tetracycline transactivator protein (tTA) under the control of the human *NEFH* promotor, and human TDP-43 (hTDP-43) with a mutated nuclear localisation sequence (ΔNLS) under the control of the *tetO* promoter (Walker et al., 2015). This allows for doxycycline (dox)-suppressible expression of cytoplasmic hTDP-43 in neurons of the brain and spinal cord, resulting in a model of ALS amenable to pre-clinical therapeutic testing (Walker et al., 2015; Spiller et al., 2019).

First, to determine if riluzole treatment could delay the characteristic onset of ALS-like motor phenotypes, we conducted a series of assessments and behavioural tests in control and rNLS mice treated daily with vehicle or riluzole for 6 weeks, beginning at the initiation of hTDP-43ΔNLS expression. Since rNLS mice exhibit weight loss during disease progression (Walker et al., 2015), we assessed the ability of riluzole to modify weight changes over time. Mice were weighed three times weekly and an average of the three weights for each week was evaluated, expressed as a percentage of starting weight on day 1. As expected, there were differences between groups over time (*F*_(15,95)_ =17.78, *p*<0.001), with significant differences between riluzole-treated control and rNLS mice from weeks 2 to 5 off dox (*p*<0.05) and vehicle-treated control and rNLS mice from weeks 3 to 5 (*p*<0.01). However, no differences were observed between vehicle-treated or riluzole-treated rNLS mice at any timepoint, indicating that riluzole treatment had no effect on weight loss in rNLS mice (Fig 1A).

**Figure 1.**
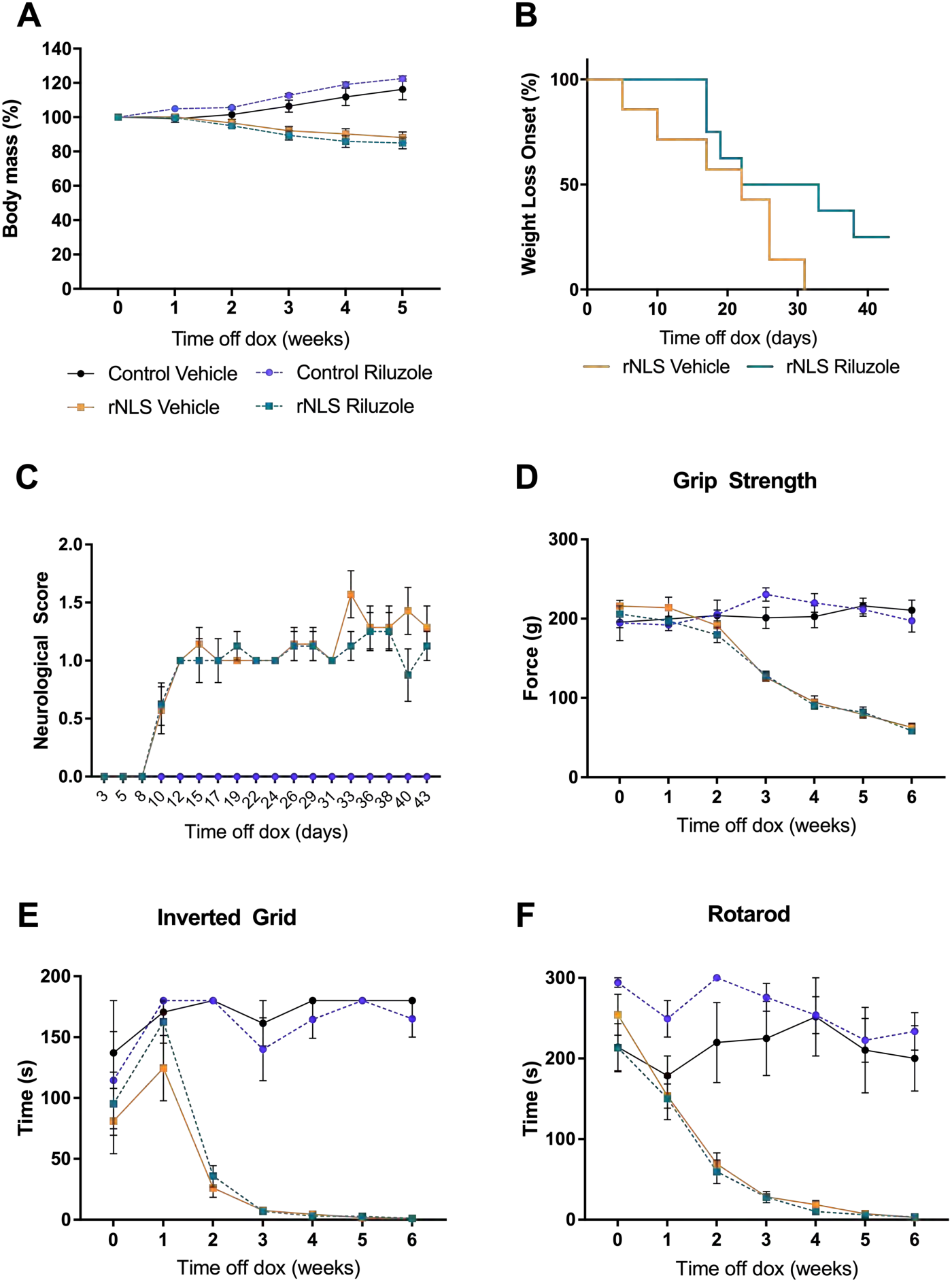
Riluzole treatment beginning at time off dox does not alter body mass change or motor behaviour in control or rNLS mice. **A** Percentage of the pre-disease (day 0) body mass showed significant weight loss over time in rNLS mice, which was not altered by riluzole. **B** Kaplan-Meier curve showing the number of days until loss of 10% of body weight. **C** No significant differences in neurological scores occurred between vehicle- or riluzole-treated rNLS mice. Behavioural assessment of motor function was performed in vehicle- or riluzole-treated control and rNLS mice from day of dox removal, for 6 weeks. No significant differences between vehicle- and riluzole-treated mice were seen in **D** grip strength, **E** inverted grid test, or **F** rotarod tests. Corresponding graph colours as per **A**, i.e. control vehicle = black solid line, control riluzole = blue dashed line, rNLS vehicle = orange solid line, rNLS riluzole = green dashed line. *n*=4/control group, *n*=7 rNLS vehicle and *n*=8 rNLS riluzole. Detailed statistical test results are described in Extended Data Table 1-1.

To assess onset of disease in vehicle- or riluzole-treated rNLS mice, we compared the number of days taken until mice reached 10% body weight loss after induction of hTDP-43ΔNLS expression. The mean number of days for vehicle-treated rNLS mice to reach a 10% body weight loss was 22 days, whereas riluzole-treated mice reached this point at a mean of 27 days, suggesting a trend towards a delay in disease onset with riluzole treatment (*p*=0.088).

We also assessed the development of neurological deficits using a 4-point scale of behavioural phenotype. We found no significant differences in the neurological scores of vehicle- or riluzole-treated rNLS mice (Fig 1C). As expected, control mice did not show signs of neurological deficits.

To assess if riluzole can improve the motor deficits observed in rNLS mice, we performed comprehensive behavioural testing. Grip strength was measured to determine changes in neuromuscular strength. As expected, dramatic muscle weakening occurred over time in rNLS mice (Fig 1D; interaction of time and group *F*_(18,114)_ = 20.66 *p*<0.0001), and a significant difference between control and rNLS groups occurred at weeks 3, 4, 5 and 6 (*p*<0.0001). There were no differences between vehicle- or riluzole-treated control mice for the full duration of the experiment, demonstrating no effect of riluzole on grip strength in wildtype animals. However, there was also no difference between vehicle- or riluzole-treated rNLS mice, indicating no effect of the treatment on grip strength in rNLS mice, despite disease-associated decline in this task over time.

To test neuromuscular balance and strength, we performed the inverted grid test on vehicle- or riluzole-treated control and rNLS mice. Again, performance for rNLS mice declined over time while control mice performed consistently throughout the experiment (Fig 1E; interaction of time and group *F*_(18,114)_ = 6.18 *p*<0.0001). Vehicle-treated rNLS mice exhibited significantly decreased time to fall from the inverted grid as compared to vehicle-treated control mice from weeks 2 to 6 (*p*<0.0001), confirming the results observed in previous studies (Walker et al., 2015). However, no differences were observed between vehicle- or riluzole-treated rNLS mice at any time point, indicating no effect of riluzole on the observed muscle weakness in rNLS mice.

As an additional test of motor impairment, we analysed the time spent on an accelerating rotarod. For rNLS mice, the time spent on the rotarod declined over time, whereas control mice performed consistently throughout the study (Fig 1F; interaction of time and group *F*_(18,114)_ = 10.23 *p*<0.0001). Similar to the grip strength and inverted grid test results, a significant difference was detected between vehicle-treated control mice and vehicle-treated rNLS mice from weeks 2 to 6 (*p*<0.0001), confirming the progressive motor deficits of the rNLS mouse model reported previously (Walker et al., 2015). However, there were no significant differences between vehicle- or riluzole-treated rNLS mice in rotarod performance, again indicating no effect of riluzole treatment on motor function in disease.

Overall, we observed no effect of riluzole on any behavioural test when mice were treated from day 0 of hTDP-43ΔNLS expression until 6 weeks after removal of dox, by which time all rNLS mice were severely impaired in motor functions.

### Riluzole does not alter soluble or insoluble TDP-43 protein levels in rNLS mice

Accumulation of aggregated and phosphorylated TDP-43 is the main pathological hallmark of ALS (Neumann et al., 2006), and thus amelioration of TDP-43 accumulation is likely to be a key marker of therapeutic efficacy in disease. Therefore, we assessed TDP-43 levels in the rNLS mice following riluzole treatment. To determine if riluzole altered the levels of soluble TDP-43 (primarily indicating non-aggregated functional protein), cortex samples from vehicle- or riluzole-treated control and rNLS mice were separated into RIPA-soluble and urea-soluble fractions (Fig 2A). Previous studies indicate robust detection of both soluble and insoluble TDP-43 in rNLS mice at 6 weeks after removal of Dox (Walker et al., 2015), and therefore this time point was chosen to assess the effects of riluzole. No differences were detected between vehicle- and riluzole-treated rNLS mice in the levels of hTDP-43ΔNLS (indicating levels of the human TDP-43 (Fig 2B). Furthermore, as expected, significant differences were detected between groups in levels of RIPA-soluble h+mTDP-43 (indicating both human and endogenous mouse TDP-43, Fig 2C; main effect of genotype *F*_(1,12)_ = 79.79 *p*<0.0001). Post-hoc analysis revealed that levels of RIPA-soluble TDP-43 did not differ in control mice regardless of riluzole treatment, although rNLS mice had significantly higher levels of RIPA-soluble TDP-43 than control mice, as expected. Notably, there were no significant differences between the levels of RIPA-soluble hTDP-43 (Fig 2B) or h+mTDP-43 (Fig 2C) in the cortex of vehicle- or riluzole-treated rNLS mice. Together these findings indicate that riluzole treatment had no effect on soluble TDP-43 levels.

**Figure 2.**
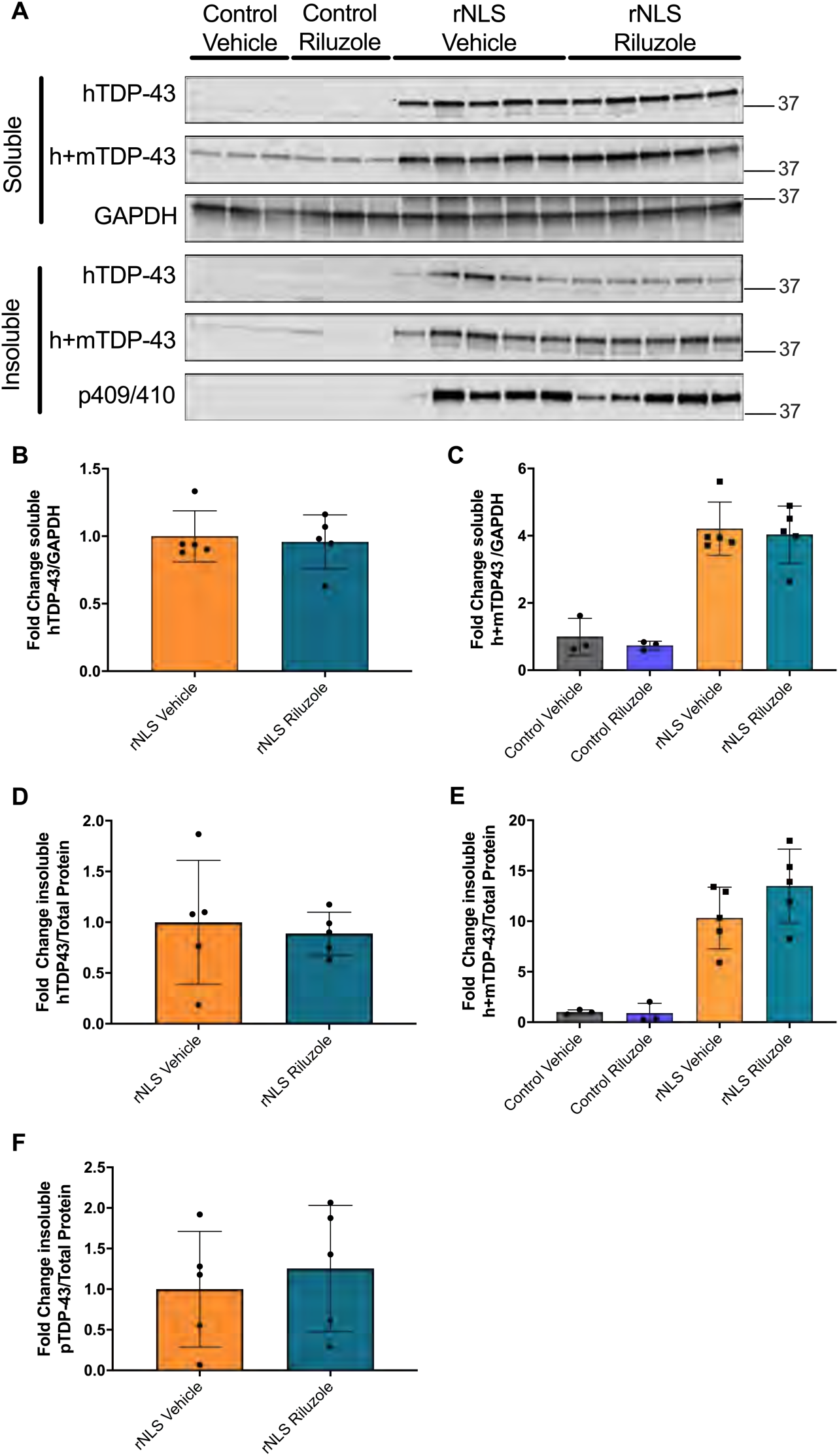
Riluzole treatment does not alter RIPA-soluble and -insoluble total and phosphorylated TDP-43 levels in control or rNLS mice. **A** Representative immunoblots of RIPA-soluble and urea-soluble cortical brain lysates from vehicle- or riluzole-treated control and rNLS mice for human-specific (h)TDP-43, human- and mouse-specific (h+m)TDP-43 and phosphorylated (p409/410) TDP-43. Quantification of **B** RIPA-soluble hTDP-43ΔNLS, **C** RIPA-soluble h+mTDP-43, **D** urea-soluble hTDP-43ΔNLS, **E** urea-soluble h+mTDP-43, and **F** p409/410 TDP-43 revealed higher levels in rNLS mice, but no effect of riluzole treatment. GAPDH for loading control of hTDP-43 and h+mTDP-43 blot is shown, with full blots and individual matched GAPDH/total protein blots used for quantification shown in Extended Data Figure 2-1. *n*=3/control group and *n*=5/rNLS group. Detailed statistical test results are described in Extended Data Table 2-1.

We next assessed levels of TDP-43 in the RIPA-insoluble/urea-soluble protein fraction, indicative of aggregated/pathological proteins. Notably, there were no statistically significant differences between vehicle- or riluzole-treated rNLS mice in urea-soluble hTDP-43ΔNLS (Fig 2D). Furthermore, despite significantly increased levels of h+mTDP-43 in rNLS mice (Fig 2E; main effect of genotype *F*_(1,12)_ = 58.33 *p*<0.0001), there was no difference between vehicle- or riluzole-treated rNLS mice, indicating that riluzole had no effect on accumulation of aggregated TDP-43 in disease. Likewise, phosphorylated TDP-43 (pS409/410) was detected only in rNLS mice in the urea-soluble fraction, and levels of phosphorylated TDP-43 were not altered between vehicle- or riluzole-treated rNLS mice (Fig 2F). Combined, these results indicate that riluzole treatment does not alter TDP-43 levels, solubility or phosphorylation in rNLS mice.

### GluA2 and GluA3, but not GluA1, glutamate receptor subunit levels are decreased in rNLS mice but are unaltered by riluzole

In the mouse brain, TDP-43 binds specifically to mRNAs of the AMPA receptor subunits *Gria2* (encoding GluA2), *Gria3* (GluA3) and *Gria4* (GluA4), but not *Gria1* (GluA1) (Sephton et al., 2011; Narayanan et al., 2013). Indeed, TDP-43 has been shown to bind to the intronic pre-mRNA of *Gria3*, and knockdown of TDP-43 in mouse brain causes both splicing changes and a decrease in levels of *Gria3* mRNA (Polymenidou et al., 2011; Lagier-Tourenne et al., 2012). However, increased surface levels of GluA1 are also observed in neurons expressing ALS-linked TDP-43^A315T^ *in vitro* (Jiang et al., 2019). Combined, these findings indicate that multiple mechanisms likely relate altered TDP-43 function to specific changes in AMPA receptor subunit levels, and these changes may contribute to excitotoxic neuron loss in ALS. Furthermore, previous work has indicated that riluzole can mediate the activity of AMPA receptors *in vitro* (Du et al., 2007), and riluzole may act in part due to non-competitive inhibition of AMPA receptors (Albo et al., 2004). Together, these studies indicate that both TDP-43 function and riluzole may affect levels and function of the glutamatergic neurotransmission machinery.

To determine if riluzole can modulate AMPA receptor levels in rNLS mice, we immunoblotted for the GluA1, GluA2 and GluA3 proteins in vehicle- or riluzole-treated control and rNLS mice (Fig 3A). There was no effect of genotype and no overall changes in GluA1 levels between each of the four groups (Fig 3B). However, levels of both GluA2 (Fig 3C; main effect of genotype *F*_(1,12)_ = 89.24 *p*<0.0001) and GluA3 (Fig 3D; main effect of genotype *F*_(1,12)_ = 55.54 *p*<0.0001) were significantly lower in rNLS mice compared to controls, suggesting specific loss of these glutamate receptor subunits during disease. However, we found no differences in GluA2 or GluA3 levels between vehicle- or riluzole-treated rNLS mice, indicating an inability of riluzole to change overall protein levels or rescue the disease-associated deficiencies of GluA2 and GluA3.

**Figure 3.**
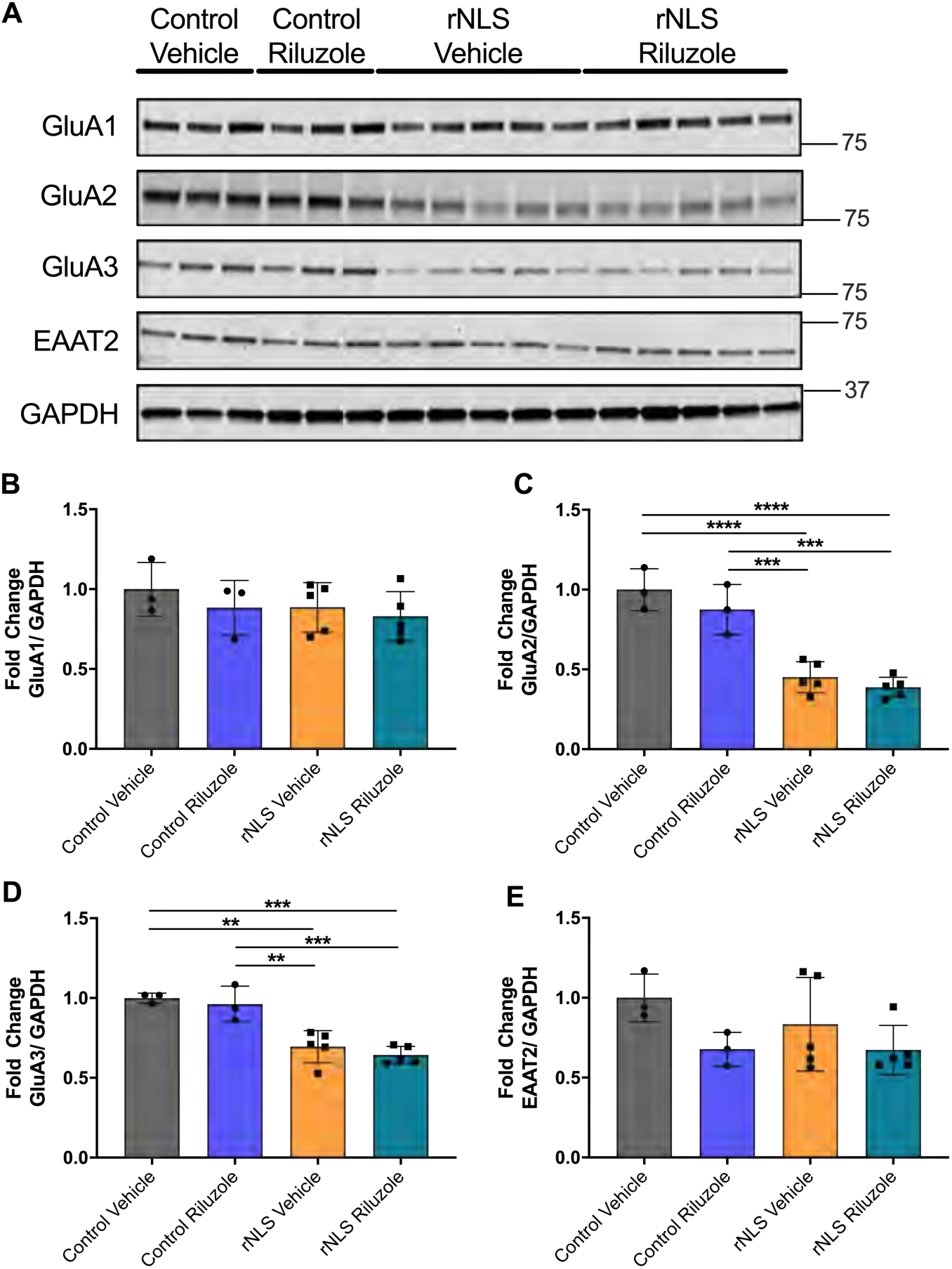
Levels of AMPA receptor subunits GluA2 and GluA3, but not GluA1 or the glutamate-transporter EAAT2, are decreased in rNLS mice but are not altered by riluzole treatment. **A** Representative immunoblots of RIPA-soluble cortical brain lysates from vehicle- or riluzole-treated control and rNLS mice for GluA1, GluA2, GluA3 and EEAT2. **B** Quantification revealed no significant differences of GluA1 between all four groups. **C** Quantification revealed significantly lower levels of GluA2 in rNLS mice compared to controls, with no effect of riluzole treatment, and **D** significantly lower levels of GluA3 in rNLS mice compared to controls, with no effect of riluzole treatment. **E** No significant changes were observed in EAAT2 levels between groups. GAPDH for loading control of GluA1 blot is shown, with full blots and individual matched GAPDH blots used for quantification shown in Extended Data Figure 3-1. *n*=3/control group and *n*=5/rNLS group. **=*p*<0.01, ***=*p*<0.001, ****=*p*<0.0001. Detailed statistical test results are described in Extended Data Table 3-1.

In addition to effects on AMPA receptor subunits, riluzole has been previously shown to upregulate expression of EAAT2 (also known as GLT-1) in astrocytes *in vitro* (Carbone et al., 2012a). We therefore assessed EAAT2 levels in vehicle- or riluzole-treated control and rNLS mice. Although there was a significant main effect of treatment (*F*_(1,12)_ = 5.244 *p*=0.0409), suggesting that riluzole may cause a small decrease in EAAT2 levels *in vivo*, there was no change to EAAT2 levels in rNLS mice compared to control mice (Fig 3E), and there were no significant differences between groups.

Overall, these results indicate a loss of glutamate receptor subunits GluA2 and GluA3 during disease, and riluzole is unable to rescue these specific deficits in rNLS mice.

### Riluzole does not prevent muscle atrophy in rNLS mice

To next address whether riluzole had an effect against disease-associated muscle atrophy, we assessed the cross-sectional area of specific muscle fibre-types in the hindlimb tibialis anterior muscle of vehicle- or riluzole-treated mice at 6 weeks following the removal of dox. We observed a large number of angulated type IIB fibres in rNLS mice (Fig 4A), indicating these fibres had been denervated (Hegedus et al., 2008). No significant differences were apparent in type IIA fibres (Figure 4B), however atrophy was observed in the type IIB fibres of rNLS mice, with approximately 60% decrease in cross-sectional area compared to controls (Fig 4C; main effect of genotype *F*_(1,14)_ = 98.46 *p*<0.0001), with no effect of riluzole treatment in rNLS mice. Type IIX muscle fibres also showed a main effect of genotype (Fig 4D; F_(1,14)_= 9.652 *p*=0.0077), suggesting type IIX muscle fibre atrophy in rNLS mice, although there were no significant differences between groups. Overall, these findings indicate that atrophy of specific muscle fibre types, in particular type IIB fibres, occurs in the rNLS mice, but this disease-associated effect is not countered by riluzole treatment.

**Figure 4.**
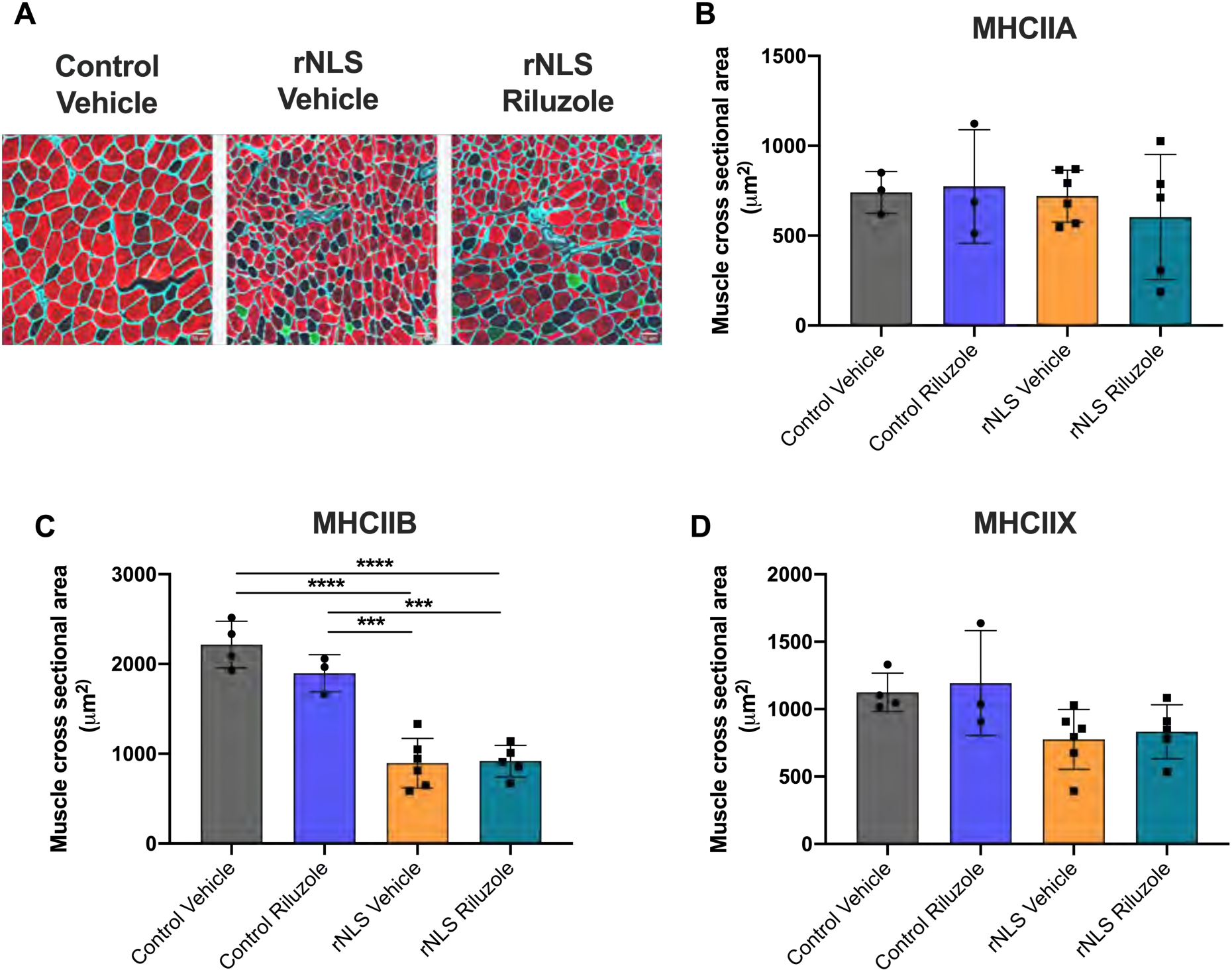
Atrophy of muscle fibres in rNLS mice, which is not affected by riluzole. **A** Representative images of tibialis anterior muscle from control vehicle, and rNLS vehicle- or riluzole-treated animals (MHCIIA, green; MHCIIB, red; laminin, cyan; MHCIIX, unstained). **B** No significant difference in type IIA muscle fibre diameter between treatment groups or genotypes. **C** rNLS mice had significantly smaller type IIB muscle fibres compared to control animals, which was not affected by riluzole. **D** No significant difference between groups occurred in type IIX muscle fibres. *n*=3/control vehicle (MHCIIA), *n*=4/control vehicle (MHCIIB, MHCIIX), n=3/control riluzole, *n*=6/rNLS vehicle and *n*=5/rNLS riluzole. \*\*\**p*<0.001, \*\*\*\**p*<0.0001. Detailed statistical test results are described in Extended Data Table 4-1.

### Riluzole does not alter developed motor deficits or extend lifespan in rNLS mice

Recent epidemiological analyses of people with ALS treated with riluzole have indicated that the disease-delaying effects of the drug are likely to occur only in the earliest and latest stages of disease (Fang et al., 2018; de Jongh et al., 2019). Since we showed above that riluzole had no effect in the earliest stages of disease in the rNLS mice, we next tested treatment beginning at 4 weeks post-disease induction, a time point when motor deficits and neurological changes had already developed in the rNLS mice (Walker et al., 2015), and studied the mice until the humane disease end-stage. We assessed mouse weights both prior to and during vehicle or riluzole treatment. Mouse body weights changed differentially over time between genotypes (Fig 5A; interaction of time and group *F*_(21,273)_ =15.44 *p*<0.0001) with decline in body weight in rNLS mice, as expected. No differences occurred between vehicle- or riluzole-treated rNLS mice either prior to or during riluzole treatment, indicating no treatment effect of riluzole in rNLS mice once weight loss was established.

**Figure 5.**
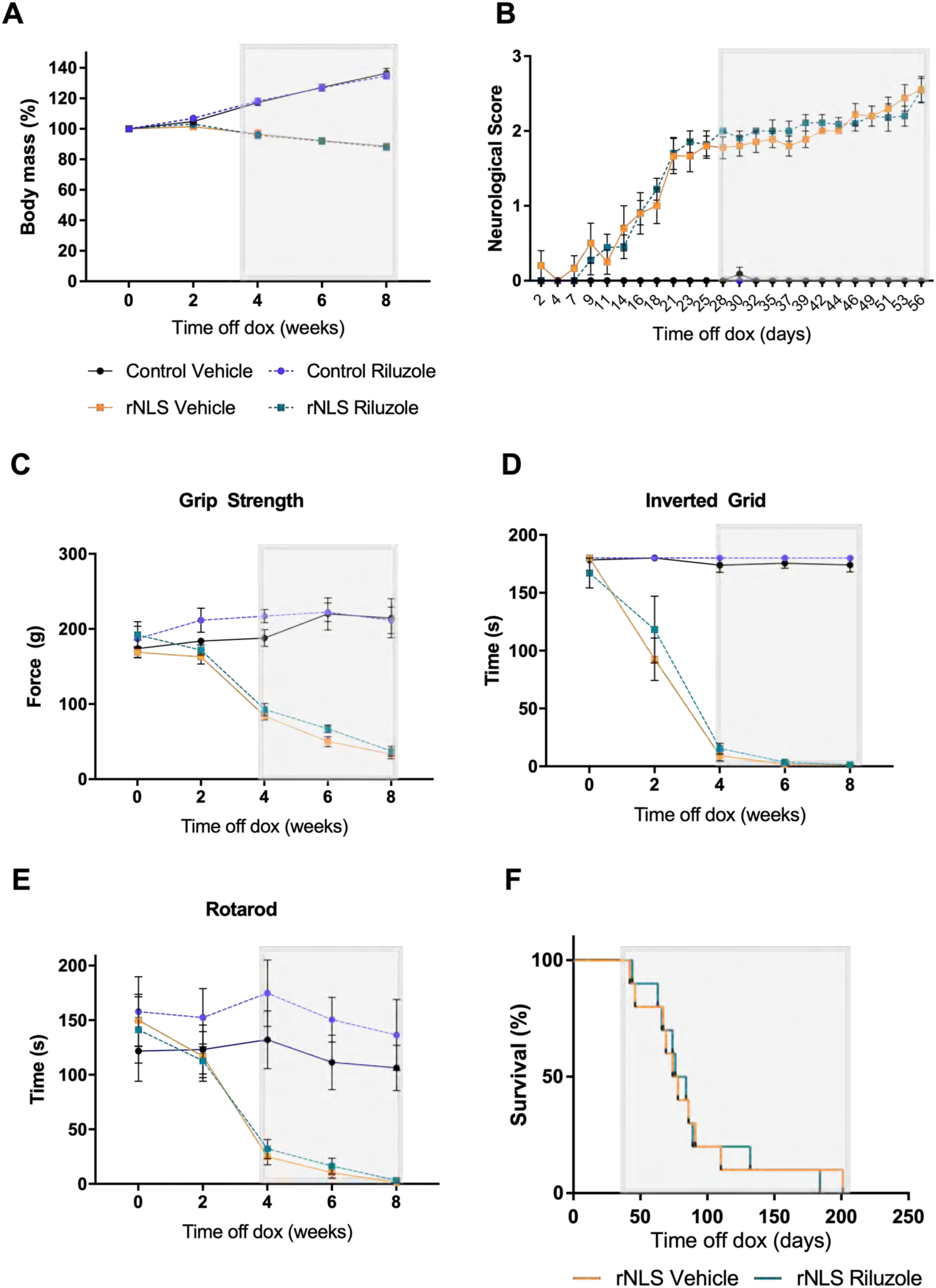
Riluzole treatment beginning after disease onset does not alter body mass loss, motor behaviour or survival in rNLS mice. **A** Percentage of the pre-disease (day 0) body mass showed significant weight loss over time in rNLS mice, which was not altered by riluzole (*n*=11 for control vehicle, control riluzole and rNLS riluzole and *n*=10 for rNLS vehicle). **B** No significant differences in neurological scores occurred between vehicle- or riluzole-treated rNLS mice (*n*=11 for control vehicle, control riluzole and rNLS riluzole and *n*=10 for rNLS vehicle). Behavioural assessment of motor function testing was analysed in a subset of vehicle- and riluzole-treated control and rNLS mice from day of dox removal, for 8 weeks. No significant differences between vehicle- or riluzole-treated mice were seen in **C** grip strength, **D** inverted grid test, or **E** rotarod tests (*n*=6/group). **F** Kaplan-Meier analysis of survival in vehicle- or riluzole-treated rNLS mice revealed no significant effect of riluzole treatment on lifespan (*n*=10/group). Corresponding graph colours as per **A**, i.e. control vehicle = black solid line, control riluzole = blue dashed line, rNLS vehicle = orange solid line, rNLS riluzole = green dashed line. Grey box indicates period during which mice were treated with vehicle or riluzole (daily from 4 weeks after removal of dox). Detailed statistical test results are described in Extended Data Table 5-1.

No signs of neurological deficits on the 4-point scale were detected in vehicle- or riluzole-treated control mice over the experimental period (Fig 5B). Neurological deficits were apparent in rNLS mice by approximately day 18. However, there were no significant differences over time between vehicle- or riluzole-treated rNLS mice (Fig 5B), indicating that riluzole treatment had no effect on development or progression of neurological deficits in the rNLS mice.

We also tested motor behaviour to determine if administration of riluzole beginning after onset of motor deficits could alter disease phenotypes. Grip strength analysis was performed both prior to and during treatment with vehicle or riluzole. Changes to muscle performance between groups occurred during the experimental period (Fig 5C; interaction of time and group *F*_(12,80)_ = 24.34 *p*<0.0001). A progressive decline in grip strength was apparent in vehicle-treated rNLS mice, with significant decline in grip strength force in rNLS mice apparent at 4, 6 and 8 weeks (*p*<0.001). However, progression of muscle weakness was not altered by post-onset treatment with riluzole, since no differences were apparent between vehicle- or riluzole-treated rNLS mice at any time point. Similar results were observed with a decline in performance of rNLS mice over time in the inverted grid test (Fig 5D; interaction of time and group *F*_(12,80)_ = 27.16 *p*<0.0001) and rotarod test (Fig 5E; interaction of time and group *F*_(12,80)_ = 7.134 *p*<0.0001). In both tests, despite significant differences between vehicle-treated control and rNLS mice at 4, 6, and 8 weeks (*p*<0.01), no changes occurred between vehicle- or riluzole-treated rNLS mice in any week, indicating that riluzole was unable to improve muscle strength and motor coordination in these animals.

Finally, we aimed to determine if riluzole could extend lifespan, since shortened survival is characteristic of rNLS mice expressing hTDP-43ΔNLS (Walker et al., 2015). Age, sex and litter-matched rNLS mice were treated with vehicle or riluzole from 4 weeks after induction of hTDP-43ΔNLS expression, and lifespan was measured from day off dox until humane disease end-stage. Analysis of the Kaplan-Meier survival curve revealed no differences between riluzole- or vehicle-treated groups (Fig 5F; χ^2^=0.01, d.f=1 and *p*=0.91), with a median survival after dox removal of 76 days for vehicle-treated and 80 days for riluzole-treated rNLS mice.

Overall, these results show that administration of riluzole starting post-disease onset (after motor deficits and neurodegeneration are already established) does not alter the behavioural phenotypes or lifespan of rNLS mice.

## Discussion

ALS is a progressive fatal neurodegenerative disease with limited therapeutic options. Riluzole was first demonstrated to provide modest survival extension to people with ALS over two decades ago and remains the most widely prescribed treatment for ALS (Lacomblez et al., 1996; Miller et al., 2012). However, the mechanisms of action of riluzole remain poorly defined and whether riluzole can modify underlying disease-related pathological changes is unclear. In this study, we sought to investigate the effects of riluzole in the well-characterised rNLS mouse model, which develops a fast and progressive ALS-like disease driven by the accumulation of cytoplasmic TDP-43 in neurons of the brain and spinal cord. We showed that riluzole administration does not alter disease onset, progression or survival of rNLS mice. Likewise, riluzole did not alter levels of soluble, insoluble or phosphorylated TDP-43. Interestingly, we found that levels of the AMPA receptor subunits GluA2 and GluA3, but not GluA1, were lower in the cortex of rNLS mice, which potentially parallels changes to glutamatergic signalling observed in people with ALS. However, riluzole treatment had no effect on this disease-related change in glutamate receptors. Likewise, we identified specific atrophy of rNLS mouse tibialis anterior muscle, in particular type MHCIIB fibres, which was not altered by riluzole treatment. Overall, our study further substantiates the limited benefit of riluzole in modifying ALS-like pathology and disease phenotypes in mice.

TDP-43 aggregation is a major neuropathological feature of ALS, and targeting of TDP-43 mislocalisation and accumulation will likely be important for development of effective disease-modifying treatments. Our study indicates that riluzole administration does not alter total levels of TDP-43 in a disease model *in vivo*, nor does riluzole affect accumulation of insoluble and phosphorylated TDP-43. Recent studies have suggested that riluzole may exert its actions through a direct link on TDP-43 self-interaction, without change to total TDP-43 levels in neuroblastoma cells *in vitro* (Oberstadt et al., 2018), although riluzole has no effect on the levels of insoluble acetylation-mimic cytoplasmic TDP-43 in cell culture (Wang et al., 2017). Furthermore, it has been hypothesised that expression of a truncated protein kinase, CK1δ, leads to phosphorylation, mislocalisation and aggregation of TDP-43 (Nonaka et al., 2016), and that riluzole is may inhibit this process (Bissaro et al., 2018). In addition, recent investigations in human iPSC-derived neurons *in vitro* demonstrated that inducing hyperexcitability was sufficient to cause TDP-43 mislocalisation (Weskamp et al., 2020). However, we observed no alteration of TDP-43 phosphorylation nor decreases in insoluble TDP-43 with riluzole treatment, indicating that riluzole may not exert direct effects against TDP-43 pathology *in vivo.*

AMPA receptors are tetrameric assemblies of four subunits designated GluA1–GluA4, which modulate fast excitatory neurotransmission in the brain. Of these, GluA2 is responsible for Ca^2+^ permeability by two mechanisms; its presence within the AMPA receptor and also by the process of RNA editing at the Q/R site by the enzyme adenosine deaminase acting on RNA 2 (ADAR2) (Wright and Vissel, 2012; Gallo et al., 2017). Indeed, a growing body of evidence indicates a deficiency in RNA editing at the GluA2 Q/R site may lead to motor neuron vulnerability in ALS (Sasaki et al., 2015; Moore et al., 2019; Yamashita and Kwak, 2019). Here, our findings of a decrease in protein levels of GluA2 and GluA3, but not GluA1, in the cortex of rNLS mice at 6 weeks after dox removal suggests that specific changes in glutamatergic neurotransmission system occur in this model during disease. Interestingly, our results provide some support for recent findings showing up-regulation of *GluA1* but decreased levels of *GluA2* and *GluA3* within the prefrontal cortex of sporadic ALS patients (Gregory et al., 2020). These alterations were not recapitulated in samples from people with a SOD1 mutation (Gregory et al., 2020), suggesting that these AMPA receptor subunit changes may be a specifically related to changes in TDP-43 function in disease.

Indeed, TDP-43-driven alteration to AMPA receptor subunits, and in particular GluA2 and GluA3, is well supported within the literature. Previous studies in mice have shown that TDP-43 binds directly to the mRNA of *Gria2* and *Gria3* (encoding GluA2 and GluA3), to regulate both splicing and total levels of *Gria3*, and TDP-43 knockdown results in lowered *Gria3* levels (Polymenidou et al., 2011; Sephton et al., 2011; Lagier-Tourenne et al., 2012; Narayanan et al., 2013). Furthermore, a loss of nuclear TDP-43 alters ADAR2 mRNA transcripts presumably leading to a reduction in GluA2 Q/R site RNA editing (Feldmeyer et al., 1999; Yamashita et al., 2013; Konen et al., 2020). Concomitantly, a loss of Q/R site RNA editing efficiency can lead to a reduction in GluA2 expression, and this may a contributing factor to loss of GluA2 observed in this study (Greger et al., 2003; Konen et al., 2020). Further research is required to determine the Q/R site RNA editing efficiency and its implications for cortical neuron loss observed in the rNLS TDP-43 model. Combined, these findings suggests that despite the dramatic accumulation of aggregation-prone cytoplasmic TDP-43 in neurons in rNLS mice (Walker et al., 2015), at least some of the molecular features that occur may be due to deregulation of TDP-43, potentially mediated by a loss of nuclear TDP-43 (Polymenidou et al., 2011). However, recent findings indicate that viral-mediated over-expression of nuclear wildtype TDP-43 in mouse brain can also cause loss of GluA1, GluA2 and GluA3 (Quadri et al., 2020), indicating an unclear relationship between TDP-43 levels and AMPA receptor subunit changes. Further investigation of how alterations in levels and localisation of AMPA receptor subunits affect disease, and how TDP-43 is involved in these processes in ALS, is warranted.

It is also worthwhile to consider other potential disease-relevant effects of riluzole for future focus of therapeutic development. Riluzole has been shown to affect persistent sodium currents (Urbani and Belluzzi, 2000; Geevasinga et al., 2016), thus altering the threshold for action potential firing, and to enhance the activity of several glutamate transporters, including the primarily astrocytic EAAT2 (also known as GLT-1) (Fumagalli et al., 2008). Riluzole also increased EAAT2 levels and activity in astrocytes in cell culture (Carbone et al., 2012b), and ameliorated the loss of EAAT2 caused by ageing or tau-associated neurodegeneration in the hippocampus of rodents (Hunsberger et al., 2015; Pereira et al., 2017). Early studies indicated that decreased glutamate transport and suppressed levels of EAAT2 in astrocytes of the brain and spinal cord are features of disease in people with ALS (Rothstein et al., 1992; Rothstein et al., 1995), and loss of EAAT2 has also been shown in post-mortem tissues in frontotemporal lobar degeneration with TDP-43 pathology (Tollervey et al., 2011). In our study, we saw no change in EAAT2 protein levels at 6 weeks after removal of dox in the rNLS mice. Likewise, we did not see an effect of riluzole on EAAT2 levels in the rNLS mice, although an effect may have been masked by the lack of a disease-related decline in total EAAT2 levels in the cortex. Astrogliosis is a feature of disease in the rNLS mice, similar to human ALS, and begins in the early stages of disease progression (Walker et al., 2015). How astrocytic changes contribute to progression of TDP-43-related disease also remains unclear, however our findings suggest that EAAT2 changes are not necessarily concomitant with astrocytic activation. Further studies are required to investigate whether the rNLS mice have alterations in intrinsic excitability or levels of glutamate transporters and other proteins involved in glutamatergic neurotransmission and glutamate scavenging, which may be protected against by riluzole, at earlier and late stages of disease. This may reveal specificity in effects and timing of these changes related to disease onset and progression. Further studies of the actions of other cell types such as microglia, which are proposed to act as important modulators of neuronal TDP-43 pathology (Spiller et al., 2018; Svahn et al., 2018), are also warranted.

In this study, we also found significant muscle atrophy in MHCIIB fibres in symptomatic rNLS mice. These results are consistent with what has been reported previously in both SOD1^G93A^ (Hegedus et al., 2008) and rNLS mice (Spiller et al., 2016). Type IIB fibres are innervated by fast-twitch fatiguable (FF) motor neurons, which are the most susceptible to dysfunction and apoptosis in ALS models, thus this fibre type is more prone to denervation, dysfunction and muscle atrophy (Nijssen et al., 2017). Likewise, in rNLS mice FF motor neuron loss occurs progressively after 4-8 weeks off dox, while fast-twitch fatigue-resistant (FR) and slow-twitch fatigue-resistant (S) motor neurons are somewhat spared (Spiller et al., 2016). However, we found that riluzole had no effect on muscle fibre cross-sectional area and thus did not protect against muscle atrophy in rNLS mice. This is consistent with human findings that revealed that riluzole has a small improvement to limb mobility, but did not improve muscle strength in people living with ALS (Miller et al., 2012).

It was previously shown that stopping the continued expression of cytoplasmic TDP-43 allowed for functional recovery in rNLS mice, even after neurodegeneration and motor impairments were evident (Walker et al., 2015). This raises hope that disease stabilisation and even improvement of symptoms is possible in people living with ALS even after disease has progressed. For treatment of ALS, it is likely that methods that modulate key aspects of disease, such as decreasing TDP-43 aggregation and mislocalisation or alterations in function and excitability of motor neurons, will need to be identified that may not individually ameliorate all aspects of disease but in combination may prove highly effective. For example, when riluzole was combined with elacridar, a drug which inhibits transporters including P-glycoprotein that control drug efflux from the central nervous system, motor function was improved and survival was significantly extended in SOD1^G93A^ mice that did not respond to riluzole alone (Jablonski et al., 2014). P-glycoprotein is up-regulated in ALS at least in part in response to astrocytic release of glutamate, and this up-regulation may decrease penetration of riluzole into the central nervous system (Mohamed et al., 2019). These findings suggest that combination therapy of other drugs with riluzole may prove to be effective, and studies of this approach in valid TDP-43 mouse models of disease are also warranted.

In summary, we have identified specific alterations in levels of some glutamatergic neurotransmission-related proteins in the rNLS mice, however riluzole, largely understood to act through this pathway, did not alter the ALS-like disease course or extend survival in this model. This is concordant with previous mouse studies that saw no effect of riluzole in SOD1^G93A^ mice (Scott et al., 2008; Hogg et al., 2018), TDP-43^A315T^ or FUS transgenic mice (Hogg et al., 2018), or TDP-43^M337V^ rats (Chen et al., 2020). These studies highlight the importance of tailoring the use of animal models to the experimental question being addressed to ensure translatability of findings, and indicate some discordance between human disease and results from widely used ALS rodent models. Indeed, the relatively rapid progression of disease in the rNLS and other ALS mouse models raises some doubt that treatments that may have at least some subtle positive effects in people can be identified in short-term animal studies. However, in future studies the rNLS model is likely to be a powerful addition for analysis of therapeutics designed to directly target dysfunctional molecular pathways related to aggregation and cytoplasmic accumulation of TDP-43, and identification of such effective therapeutics offer the best chances of translation. Our studies reported here demonstrate that the rNLS mice have additional disease-reminiscent features not previously identified, including specific loss of AMPA receptor subunits and atrophy of MHCIIB muscle fibres, which may also be useful markers in future pre-clinical studies of disease-targeted therapies. Renewed focus on understanding the mechanisms of TDP-43 pathology formation and other core molecular signatures of ALS, and the design of new strategies to modulate the upstream causes of disease, offer potential for more effective treatments for ALS and other devastating neurodegenerative diseases.

## Author Contributions

ALW, PADG, SL and AKW Designed research; ALW, PADG, SL, BAB, PM, KRJ, HG, RSG, TJH, WRR, OW, JV and AKW Performed research; RSC, JDA, AL, BS, CAB and MM Contributed unpublished reagents/analytic tools; ALW, PADG, SL, SSK and AKW Analyzed data; ALW, PADG, SL, CAB, MM and AKW Wrote the paper.

## Acknowledgements

We thank Prof. Virginia Lee (University of Pennsylvania) for the TDP-43 antibody (clone 5104) and Christine Sutter and the staff of the Central Animal Facility at Macquarie University for animal husbandry.

## Funding sources

This work was supported by an Australian Government Research Training Program Scholarship to BAB, the MND Research Institute of Australia (PhD top-up scholarship to BAB and Cure for MND Research Grant 1651 to AKW), the Australian National Health and Medical Research Council (Project Grant 1124005 and RD Wright Career Development Fellowship 1140386 to AKW), donations made to the Macquarie University Centre for Motor Neuron Disease Research, the Ross Maclean Fellowship, and the Brazil Family Program for Neurology.

**Extended Data Figure 2-1.**
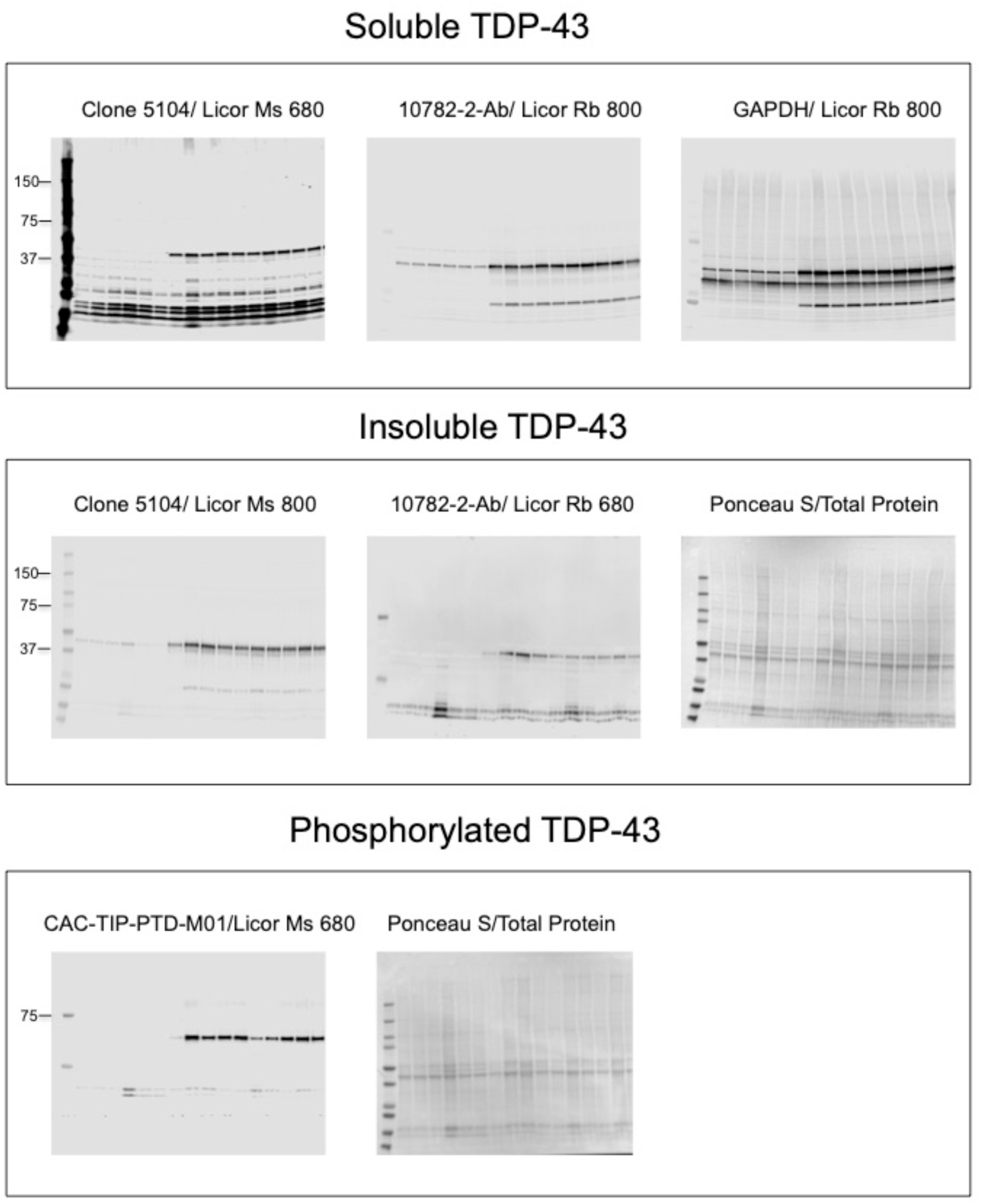
Full immunoblots for TDP-43 levels from Figure 2 and matched associated loading control blots used for protein quantification. For soluble TDP-43, GAPDH was used for normalisation. For insoluble fractions (TDP-43 and p-TDP-43), protein levels were normalised against full-length Ponceau S lanes.

**Extended Data Figure 3-1.**
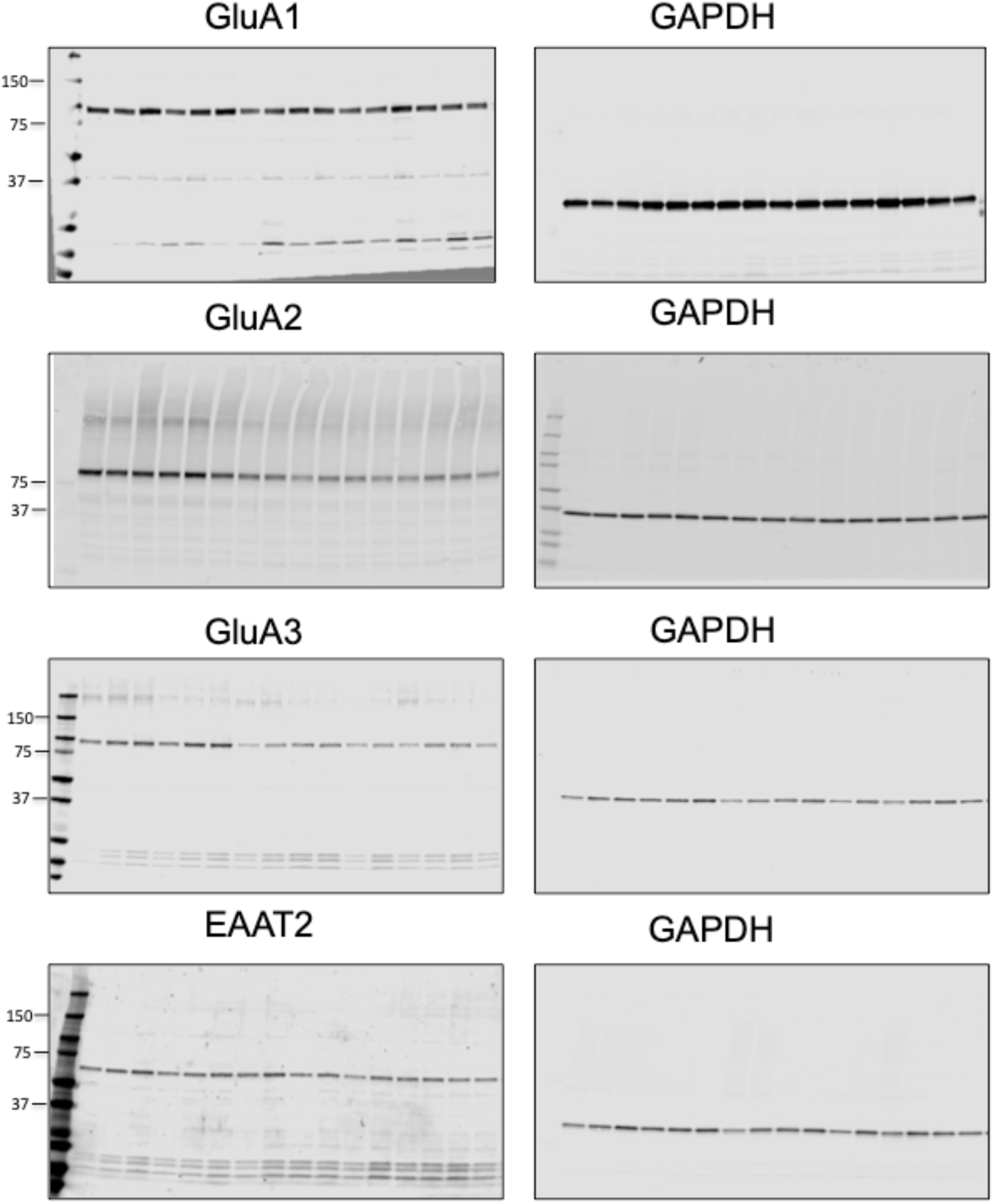
Full immunoblots for glutamate receptor subunit and glutamate transporter protein levels from Figure 3, and matched associated GAPDH blots used for protein quantification.

**Extended Data Table 1-1.** Description of statistical results of all tests shown in Figure 1.

| Analysis | Statistical Test | Main Effect | F (DFn, DFd) | P value |
| --- | --- | --- | --- | --- |
| Fig 1A. Body Weight | RM Two-Way ANOVA | Time x Group | F (15, 95) = 17.48 | P<0.0001 |
|  |  | Time | F (5, 95) = 1.701 | P=0.1418 |
|  |  | Group | F (3, 19) = 20.28 | P<0.0001 |
|  |  | Subject | F (19, 95) = 6.326 | P<0.0001 |
| Fig 1B. Days till 10% Onset | Log-rank (Mantel-Cox) | Chi-square=2.918, d.f 1 | N/A | 0.0876 |
| Fig 1C. NS by day | Mixed Model | Time | F (17, 322) = 15.79 | <0.0001 |
|  |  | Group | F (3, 19) = 67.39 | <0.0001 |
|  |  | Time x Group | F (51, 322) = 5.780 | <0.0001 |
| Fig 1D. Grip Strength | RM Two-Way ANOVA | Time x Group | F (18, 114) = 20.66 | <0.0001 |
|  |  | Time | F (6, 114) = 41.94 | <0.0001 |
|  |  | Group | F (3, 19) = 49.13 | <0.0001 |
|  |  | Subject | F (19, 114) = 2.871 | 0.0003 |
| Fig 1E. Inverted Grid | RM Two-Way ANOVA | Time x Group | F (18, 114) = 6.179 | <0.0001 |
|  |  | Time | F (6, 114) = 12.75 | <0.0001 |
|  |  | Group | F (3, 19) = 80.08 | <0.0001 |
|  |  | Subject | F (19, 114) = 2.001 | 0.0133 |
| Fig 1F. Rotarod | RM Two-Way ANOVA | Time x Group | F (18, 114) = 10.23 | <0.0001 |
|  |  | Time | F (6, 114) = 33.82 | <0.0001 |
|  |  | Group | F (3, 19) = 37.02 | <0.0001 |
|  |  | Subject | F (19, 114) = 6.846 | <0.0001 |

**Extended Data Table 2-1.** Description of statistical results of all tests shown in Figure 2.

| Analysis | Statistical Test | Main Effect | t, df / F (DFn, DFd) | P value |
| --- | --- | --- | --- | --- |
| Fig 2B. hTDP-43 soluble | Unpaired t-test | N/A | t=0.3016, df=8 | 0.7707 |
| Fig 2C. h+m TDP-43 soluble | Two-Way ANOVA | Interaction | F (1, 12) = 0.01352 | 0.9094 |
|  |  | Genotype | F (1, 12) = 79.79 | <0.0001 |
|  |  | Treatment | F (1, 12) = 0.3639 | 0.5576 |
| Fig 2D. hTDP-43 insoluble | Unpaired t-test | N/A | t=0.3863, df=8 | 0.7093 |
| Fig 2E. h+m TDP-43 insoluble | Two-Way ANOVA | Interaction | F (1, 12) = 1.293 | 0.2778 |
|  |  | Genotype | F (1, 12) = 58.33 | <0.0001 |
|  |  | Treatment | F (1, 12) = 1.152 | 0.3043 |
| Fig 2F. pTDP-43 insoluble | Unpaired t-test | N/A | t=0.5410, df=8 | 0.6032 |

**Extended Data Table 3-1.**
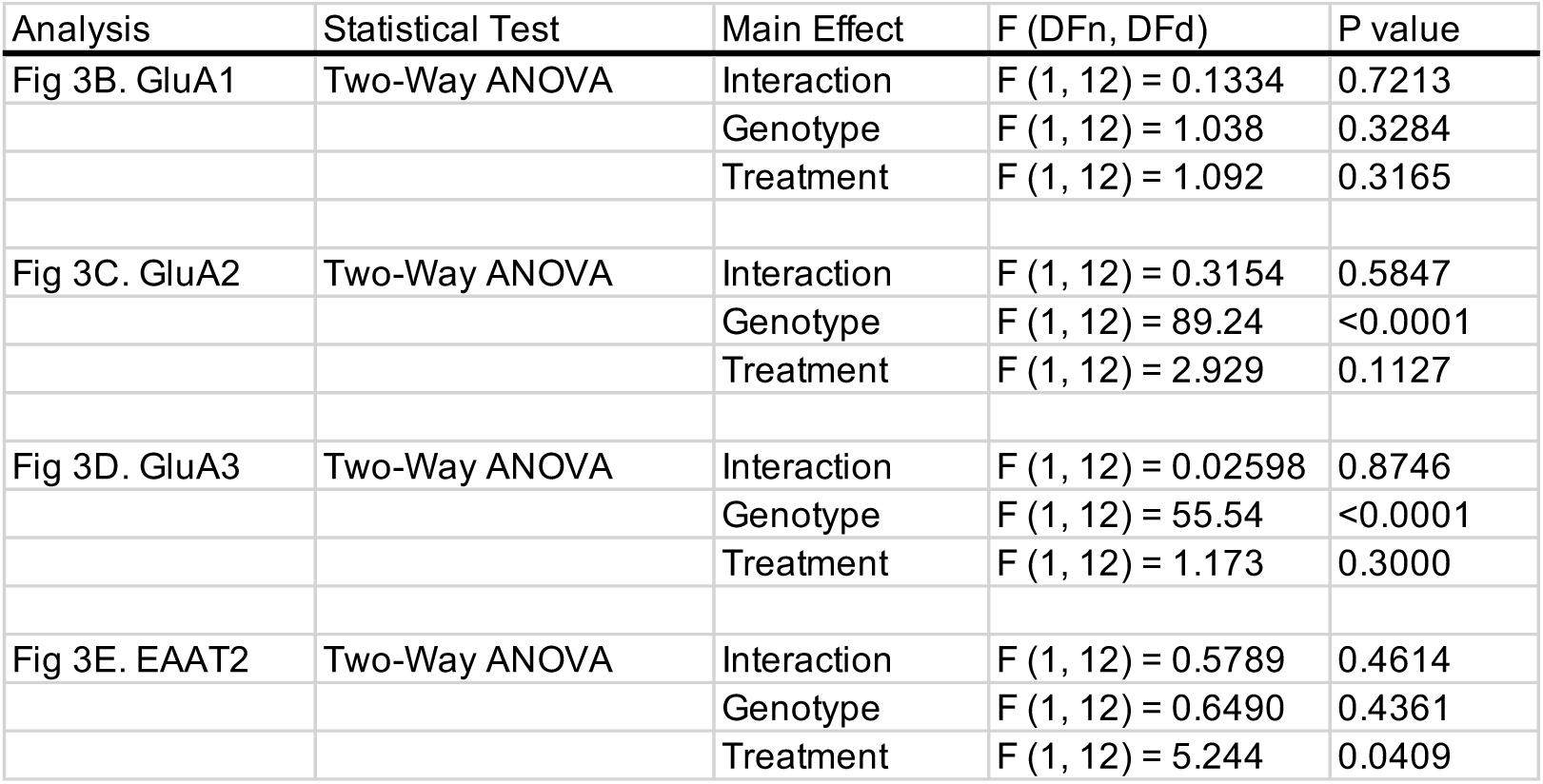
Description of statistical results of all tests shown in Figure 3.

| Analysis | Statistical Test | Main Effect | F (DFn, DFd) | P value |
| --- | --- | --- | --- | --- |
| Fig 3B. GluA1 | Two-Way ANOVA | Interaction | F (1, 12) = 0.1334 | 0.7213 |
|  |  | Genotype | F (1, 12) = 1.038 | 0.3284 |
|  |  | Treatment | F (1, 12) = 1.092 | 0.3165 |
| Fig 3C. GluA2 | Two-Way ANOVA | Interaction | F (1, 12) = 0.3154 | 0.5847 |
|  |  | Genotype | F (1, 12) = 89.24 | <0.0001 |
|  |  | Treatment | F (1, 12) = 2.929 | 0.1127 |
| Fig 3D. GluA3 | Two-Way ANOVA | Interaction | F (1, 12) = 0.02598 | 0.8746 |
|  |  | Genotype | F (1, 12) = 55.54 | <0.0001 |
|  |  | Treatment | F (1, 12) = 1.173 | 0.3000 |
| Fig 3E. EAAT2 | Two-Way ANOVA | Interaction | F (1, 12) = 0.5789 | 0.4614 |
|  |  | Genotype | F (1, 12) = 0.6490 | 0.4361 |
|  |  | Treatment | F (1, 12) = 5.244 | 0.0409 |

**Extended Data Table 4-1.** Description of statistical results of all tests shown in Figure 4.

| Analysis | Statistical Test | Main Effect | F (DFn, DFd) | P value |
| --- | --- | --- | --- | --- |
| Fig 4B. IIA | Two-Way ANOVA | Interaction | F (1, 13) = 0.3504 | 0.564 |
|  |  | Genotype | F (1, 13) = 0.5643 | 0.4659 |
|  |  | Treatment | F (1, 13) = 0.1100 | 0.7454 |
| Fig 4C. IIB | Two-Way ANOVA | Interaction | F (1, 14) = 2.212 | 0.1591 |
|  |  | Genotype | F (1, 14) = 98.46 | <0.0001 |
|  |  | Treatment | F (1, 14) = 1.675 | 0.2166 |
| Fig 4D. IIX | Two-Way ANOVA | Interaction | F (1, 14) = 0.002898 | 0.9578 |
|  |  | Genotype | F (1, 14) = 9.652 | 0.0077 |
|  |  | Treatment | F (1, 14) = 0.2978 | 0.5938 |

**Extended Data Table 5-1.** Description of statistical results of all tests shown in Figure 5.

| Analysis | Statistical Test | Main Effect | F (DFn, DFd) | P value |
| --- | --- | --- | --- | --- |
| Fig 5A. Body Weight | RM Two-Way ANOVA | Time x Group | F (21, 273) = 15.44 | <0.0001 |
|  |  | Time | F (7, 273) = 71.32 | <0.0001 |
|  |  | Group | F (3, 39) = 12.22 | <0.0001 |
|  |  | Subject | F (39, 273) = 18.37 | <0.0001 |
| Fig 5B. NS by day | Mixed Model | Time | F (23, 782) = 71.75 | <0.0001 |
|  |  | Group | F (3, 39) = 234.5 | <0.0001 |
|  |  | Time x Group | F (69, 782) = 24.37 | <0.0001 |
| Fig 5C. Grip Strength | RM Two-Way ANOVA | Time x Group | F (12, 80) = 24.34 | <0.0001 |
|  |  | Time | F (4, 80) = 30.60 | <0.0001 |
|  |  | Group | F (3, 20) = 35.14 | <0.0001 |
|  |  | Subject | F (20, 80) = 5.188 | <0.0001 |
| Fig 5D. Inverted Grid | RM Two-Way ANOVA | Time x Group | F (12, 80) = 27.16 | <0.0001 |
|  |  | Time | F (4, 80) = 84.86 | <0.0001 |
|  |  | Group | F (3, 20) = 295.8 | <0.0001 |
|  |  | Subject | F (20, 80) = 1.103 | 0.3633 |
| Fig 5E. Rotarod | RM Two-Way ANOVA | Time x Group | F (12, 80) = 7.134 | <0.0001 |
|  |  | Time | F (4, 80) = 26.23 | <0.0001 |
|  |  | Group | F (3, 20) = 6.383 | 0.0033 |
|  |  | Subject | F (20, 80) = 9.097 | <0.0001 |
| Fig 5F. Survival | Log-rank (Mantel-Cox) | Chi-square=0.0118, d.f 1 | N/A | 0.913 |

